# Mechanistic Insights into ADAR1-p110 Nuclear Import: A Dual-Interface Interaction with Transportin-1

**DOI:** 10.64898/2026.11.17.767233

**Authors:** Danielle Huey Ren Chin, Jesslyn Zhou, Yongbo Luo, Davin Hong, Yaw Bia Tan, Polly Leilei Chen, Dahai Luo

## Abstract

The human ADAR1 exists as two isoforms: the predominantly cytoplasmic-localized p150, recognized for its role in suppression of aberrant immune activation, and the predominantly nuclear-localized p110 (missing the N terminal Zα domain), whose biological functions remain less understood but are thought to rely heavily on its subcellular localization. Despite its structural similarity to the Zα domain unique to ADAR1-p150, the functional significance of the Zβ domain shared by both isoforms remains unclear. Here, we identify an additional mechanism contributing to the nuclear import of ADAR1-p110 through a dual-interface interaction with transportin-1 (TNPO1), involving both the Zβ and dsRBD3 domains of ADAR1. Using cryo-electron microscopy, we show that Zβ engages TNPO1 at its canonical cargo-binding groove, while dsRBD3 binds at a peripheral site overlapping the RanGTP binding interface. Structure-guided mutagenesis and biochemical analyses support the contribution of Zβ to TNPO1 recognition. Disruption of the interaction between ADAR1 Zβ domain and TNPO1 alters the nucleocytoplasmic distribution of ADAR1-p110, resulting in increased cytoplasmic accumulation despite the retention of the previously characterized dsRBD3-associated nuclear localization signal. Furthermore, cytoplasmic redistribution of ADAR1-p110 mutants is accompanied by global suppression of cytosolic innate immune signaling pathways and reduced expression of interferon-stimulated genes. Together, these findings identify ADAR1 Zβ as an additional TNPO1-recognition element synergizing with dsRBD3 that contributes to ADAR1-p110 nuclear localization and highlight the relationship between its subcellular distribution and immune regulatory activity.

**Significance Statement:** The subcellular localization of adenosine deaminase acting on RNA 1 (ADAR1) is critical for its biological functions, particularly in regulating innate immune responses. However, the mechanisms governing nuclear import of its p110 isoform remain incompletely understood. Here, we uncover an additional nuclear import mechanism mediated by the previously uncharacterized Zβ domain, which collaborates with dsRBD3 to engage transportin-1 (TNPO1) through two distinct binding interfaces. Disrupting Zβ-mediated recognition alters ADAR1-p110 nucleocytoplasmic distribution and is associated with suppression of cytosolic innate immune signalling and interferon-stimulated gene expression. These findings establish a previously unrecognized function of Zβ in ADAR1 trafficking and reveal a connection between ADAR1-p110 localization and immune regulation.

## Introduction

ADAR1 is an RNA-editing enzyme that catalyzes the deamination of adenosine to inosine (A-to-I) in double-stranded RNA (dsRNA) substrates [1, 2]. Due to inosine’s structural resemblance to guanosine, such edits can influence RNA stability, gene regulation, coding potential, and innate immune recognition [3]. Loss or mutations in ADAR1 typically result in aberrant sensing of endogenous double-stranded RNA and inappropriate activation of innate immune sensors [4–6]. The ADAR1 gene is expressed as two major isoforms: the interferon-inducible, predominantly cytoplasmic p150 and the constitutively expressed, predominantly nuclear p110 [7, 8]. While the p150 isoform has been widely studied for its role in cytoplasmic immune regulation [9], the biological roles of the ADAR1-p110 beyond nuclear dsRNA editing remain poorly understood. A recent study using p110 isoform-specific mouse knock-out models demonstrated that ADAR1-p110 plays an essential role in postnatal survival via mechanisms that are both editing- and innate immune-independent [10], suggesting that p110 has essential functions beyond RNA editing. Indeed, ADAR1-p110 has been implicated in various aspects of RNA metabolism, including regulation of 3’-untranslated regions [11], modulation of RNA interference pathways [12], alternative splicing mechanisms [13], and regulation of R-loop homeostasis [14, 15]. Under stress conditions, ADAR1-p110 phosphorylation has been linked to cytoplasmic export, where it prevents apoptosis by stabilizing anti-apoptotic transcripts [16]. These studies highlight the distinct biology of p110 from p150, with several reported functions dependent on its precise subcellular localization. Such regulated nucleocytoplasmic shuttling emphasizes how the precise localization of p110 is critical for cellular homeostasis. Inappropriate cytoplasmic accumulation could alter its accessibility to cytoplasmic RNA substrates and immune regulatory pathways, although the functional consequences of such redistribution remain incompletely understood.

Nuclear import of ADAR1 has been attributed to a bipartite, non-canonical nuclear localization signal (NLS) embedded in the flanking regions of dsRBD3 that interact with the import protein TNPO1. Specifically, the solution structure of ADAR1-dsRBD3 revealed that this atypical NLS is assembled within an α-helix extension adjoining the dsRBD core [17, 18]. Functional studies have shown that these flanking regions are both necessary and sufficient for TNPO1-dependent nuclear import [17]. Notably, nuclear import is further tuned by RNA binding which disrupts the dsRBD3-TNPO1 interaction, coupling nuclear import competency to the enzyme’s RNA-binding state [17], ensuring that unbound, transcriptionally “ready” ADAR1 is imported into the nucleus, while RNA-bound forms remain localized in the cytoplasm. However, proteomic profiling of ADAR1-interacting proteins suggests additional regulatory complexity beyond this model, with distinct nuclear and cytoplasmic interactomes implying other mechanisms of localization control [19, 20]. Notably, a previous study identified putative nuclear localization signals within the N-terminal regions of human and rat ADARs [21], suggesting the presence of additional localization signals.

The Zβ domain, present in both ADAR1 isoforms, is structurally conserved and adopts an αβααββα topology composed of four α-helices and three β-strands, placing it within the helix-turn-helix (HTH) family of Z-DNA binding proteins [22]. Despite its high structural similarity to Zα, several residues implicated in Zα-mediated Z-form nucleic acid recognition, suggesting differences in their nucleic acid binding properties [22, 23]. Recent biochemical studies have demonstrated that Zβ selectively recognizes RNA and DNA G-quadruplexes with nanomolar affinities, although the physiological significance of this interaction remains unclear [24]. Given its conservation and its structural divergence from the Zα nucleic acid binding interface, Zβ has been hypothesized to serve as a protein–protein interaction module. In light of emerging evidence that structured domains beyond classical NLS can mediate TNPO1 binding [17, 25–29], we posited that Zβ may act as an additional TNPO1-recognition element alongside the previously characterized dsRBD3-associated NLS.

To address how ADAR1-p110 is imported into the nucleus and how this process is coupled to its immune regulatory functions, we determined the cryo-EM structure of TNPO1 bound to the Zβ and dsRBD3 domains of ADAR1. The structure revealed that Zβ engages the canonical cargo-binding groove of TNPO1, whereas dsRBD3 binds a distinct non-canonical site on the N-terminal region. Guided by this model, we used mutagenesis and functional assays to dissect how these interactions contribute to the regulation of the subcellular localization of ADAR1-p110 and influence its ability to modulate innate immune signaling.

## Results

### Cryo-EM structure of ADAR1 binding to TNPO1

To determine how ADAR1 is recognized by its nuclear import factor - TNPO1, we solved the cryo-EM structure of TNPO1 in complex with the ADAR1 (aa 128 – aa 812) which include the Za-Zβ-dsRBD1-2-3 domains of ADAR1 (Fig. 1a). Using size exclusion chromatography, we observed a shift in the elution profile of the TNPO1-ADAR1 complex relative to the individual proteins (Fig. 1b), consistent with complex formation in solution. Single-particle cryo-EM reconstruction yielded a structure of the complex at an overall resolution of 3.15 Å (Fig. S1, Table S1). While TNPO1 was resolved across much of the reconstructed density, only the Zβ domain (residues L294–N370) and dsRBD3 (residues K715–F804) were traceable within ADAR1 (Fig. 1c-d), consistent with their engagement with TNPO1 and the likely flexibility of the interdomain-regions of ADAR1.

**Figure 1.**
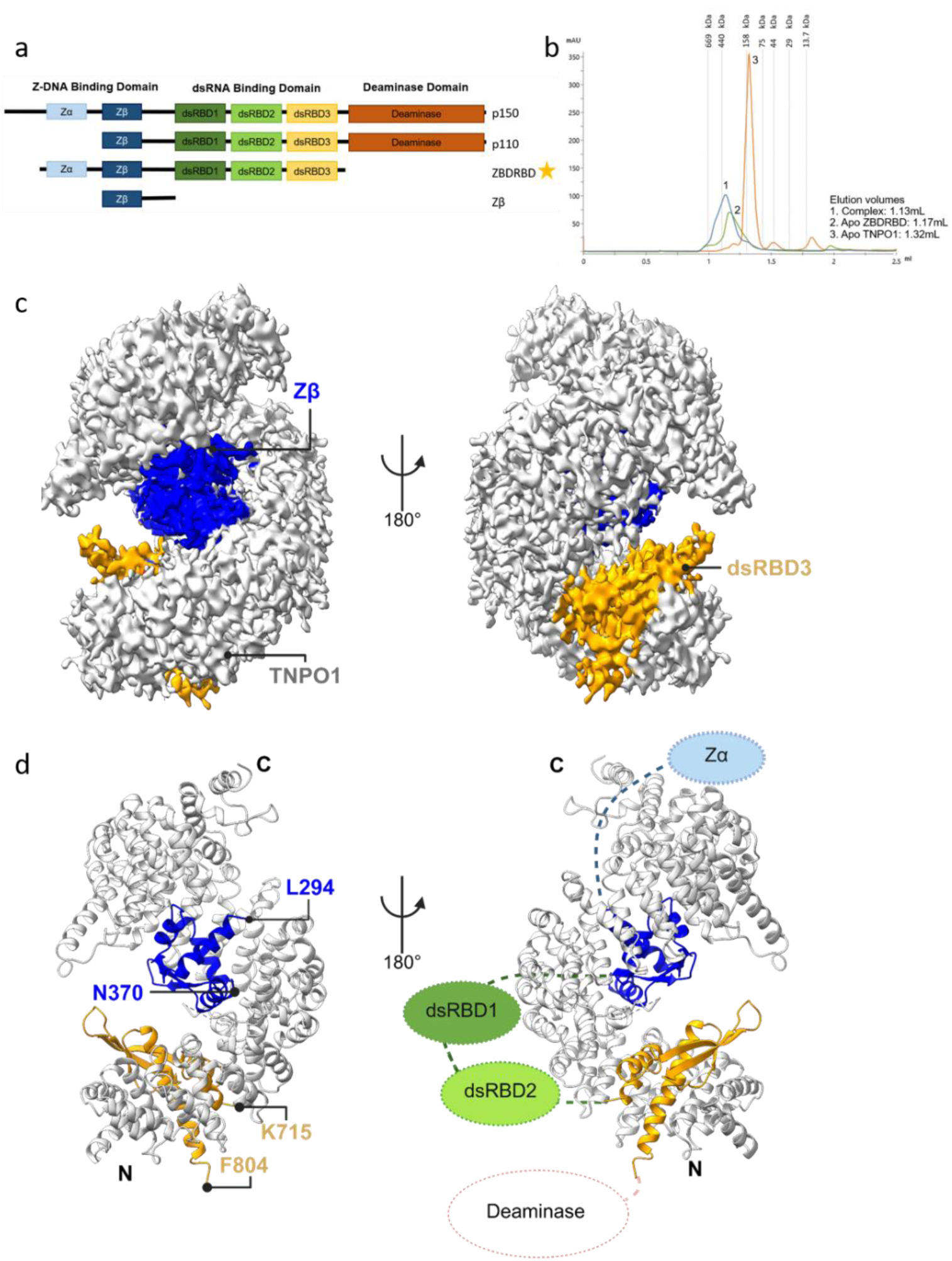
Cryo-EM structure of ADAR1 binding to TNPO1. **a.** Schematic diagram of ADAR1 constructs. Construct used for cryo-EM complex is starred. Zα: Z-NA binding domain alpha, Zβ: Z-NA binding domain beta, dsRBD: double-stranded RNA binding domain. **b.** Comparison chromatogram of apo-TNPO1 (orange; 128.5 kDa), apo-ZBD-RBD (green; 79.7 kDa) and TNPO1 complexed with ADAR1 ZBDRBD construct (blue). Analytical gel filtration performed on S200 increase 3.2/300 column. Void volume is approximately 0.8 mL. Cryo-EM **c.** map and **d.** model of the TNPO1-ZBDRBD complex. Full-length TNPO1 colored in grey, ADAR1 Zβ domain (L294-N370) and dsRBD3 domain (K715-F804) colored in blue and yellow respectively. A schematic diagram of the ADAR1-TNPO1 complex is prepared where the missing domains of ADAR1 ZBDRBD are coloured and filled using dotted lines.

To further evaluate these domain assignments, we compared the fitting of the structurally related ADAR1 domains into their respective cryo-EM densities. Comparison of the Zα and Zβ domains revealed regions of poorer correspondence between the Zα model and the observed density, supporting the assignment of Zβ in this position (Fig. S2). Similarly, comparisons of dsRBD1, dsRBD2 and dsRBD3 identified regions of poorer density correspondence for the alternative dsRBD models, particularly at the extended NLS helices, supporting the assignment of dsRBD3 (Fig. S3). These comparisons provide additional structural support for the proposed identities of the two TNPO1-associated ADAR1 domains.

The cryo-EM map and model revealed that Zβ adopts a compact, globular fold that anchors into the central cavity of the TNPO1 superhelical solenoid—a site typically occupied by classical PY-NLS cargoes. In contrast, dsRBD3 binds to a more peripheral region near the N terminus of TNPO1, overlapping with the known RanGTP binding site (Fig. 1c-d). Structural comparisons with the crystallographic and NMR structures showed that both Zβ and dsRBD3 adopt conformations closely matching their isolated forms, with root-mean-square deviations (RMSDs) of 1.00 Å (Zβ; 60 aligned Cα atoms, reference PDB: 1XMK [22]) (Fig. S4a) and 0.91 Å (dsRBD3; 73 aligned Cα atoms, reference PDB: 2MDR [16]) (Fig. S4b), respectively. While TNPO1 remains globally well-structured, the overall RMSD of 4.98 Å across 808 aligned Cα atoms suggests conformational changes, particularly toward the C-terminal end of the solenoid (Fig. S4c). These differences may reflect structural accommodation of the insertion of Zβ into the central cargo-binding groove.

Taken together, the cryo-EM structure supports a composite TNPO1–ADAR1 interface in which Zβ and dsRBD3 engage distinct regions of the TNPO1, providing a structural framework for a proposed dual-domain recognition mechanism that contributes to ADAR1-p110 nuclear import.

### TNPO1 uses both canonical and extended binding interfaces to bind ADAR1

Closer examination of the TNPO1–ADAR1 complex revealed that the Zβ and dsRBD3 domains contact TNPO1 through spatially distinct surfaces. The Zβ domain engages the canonical cargo-binding groove formed along the inner concave surface of the TNPO1 solenoid (Fig. 1d, 2a, 2c). This interface is characterized by shape complementarity and an extensive network of polar and electrostatic interactions, consistent with known features of classical importin–cargo recognition. Zβ presents a positively charged surface composed primarily of residues K320, R328, R345, K358, R362, and K366, along with the polar residues N317 and N331, which collectively form a broad electrostatic interface that aligns with the negatively charged cargo pocket of TNPO1. These residues participate in multiple hydrogen bonds with conserved acidic residues on TNPO1 (Fig. 2c)—such as E286 and D647—that are also utilized in the recognition of classical PY-NLS motifs. Quantitative analysis revealed that the Zβ–TNPO1 interface buries approximately 1,356 Å² of solvent-accessible surface area (Table S2).

**Figure 2.**
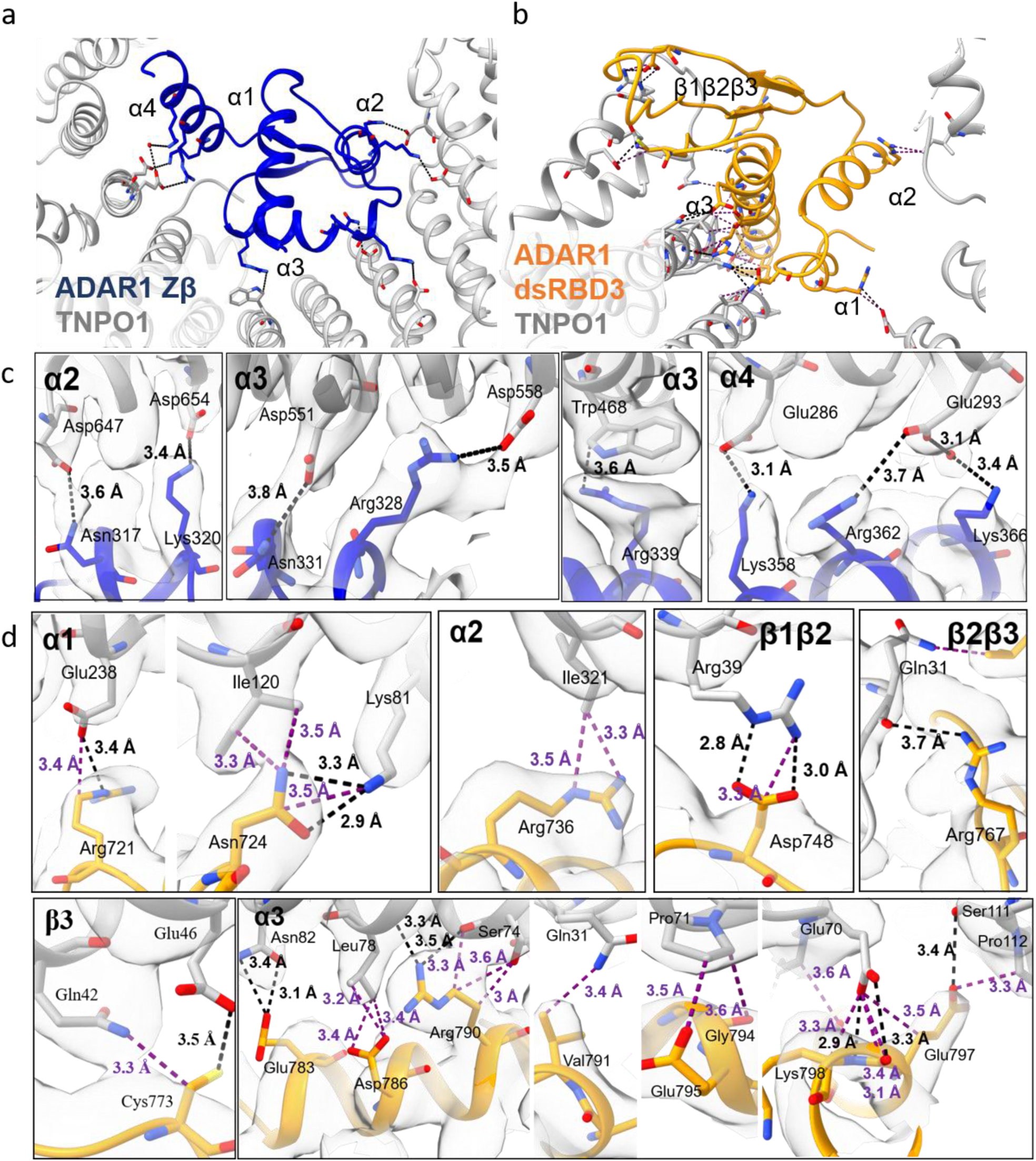
ADAR1 forms a dual-domain recognition mechanism to engage TNPO1. Model structure of **a.** ADAR1-Zβ with TNPO1 and **b.** ADAR1-dsRBDIII and TNPO1. Local side chain fitting of **c.** Zβ and **d.** dsRBD3 with TNPO1 at the residue level. All interactions are labelled with the corresponding bond lengths. Electrostatic/hydrogen bonds are depicted in black; van der Waals interactions are depicted in purple. TNPO1 colored in grey, and ADAR1 Zβ and dsRBD3 domain is coloured in blue and yellow respectively.

Structural alignment with known TNPO1–cargo complexes (Fig. 3a-b), including those containing PY-NLS peptides (PDB entries 2OT8 [26], 5TQC [30], 8SGH [31], 4OO6 [32], 4FDD [33], and 5YVH [34]), revealed that Zβ partially recapitulates the spatial arrangement and electrostatic surface potential characteristic of canonical motifs. However, Zβ lacks the hallmark proline–tyrosine (PY) dipeptide (Fig. 3b, 4a) and does not meet all criteria typically used to define this class of signals. Comparative interface analysis revealed that the examined PY-NLS–TNPO1 complexes exhibit buried surface areas ranging from approximately 444 Å^2^ to 1,098 Å^2^ (Table S2), all of which were consistently smaller than that of Zβ-TNPO1. Notably, the FUS–TNPO1 complexes (PDB: 4FDD and 5YVH), which contain longer PY-NLS constructs, exhibited buried surface areas of approximately 1,098 Å^2^ and 1,014 Å^2^, respectively (Table S2). The Zβ–TNPO1 interface is therefore approximately 24–34% larger than these two representative complexes, despite differences in the structural organization of the interacting cargoes.

**Figure 3.**
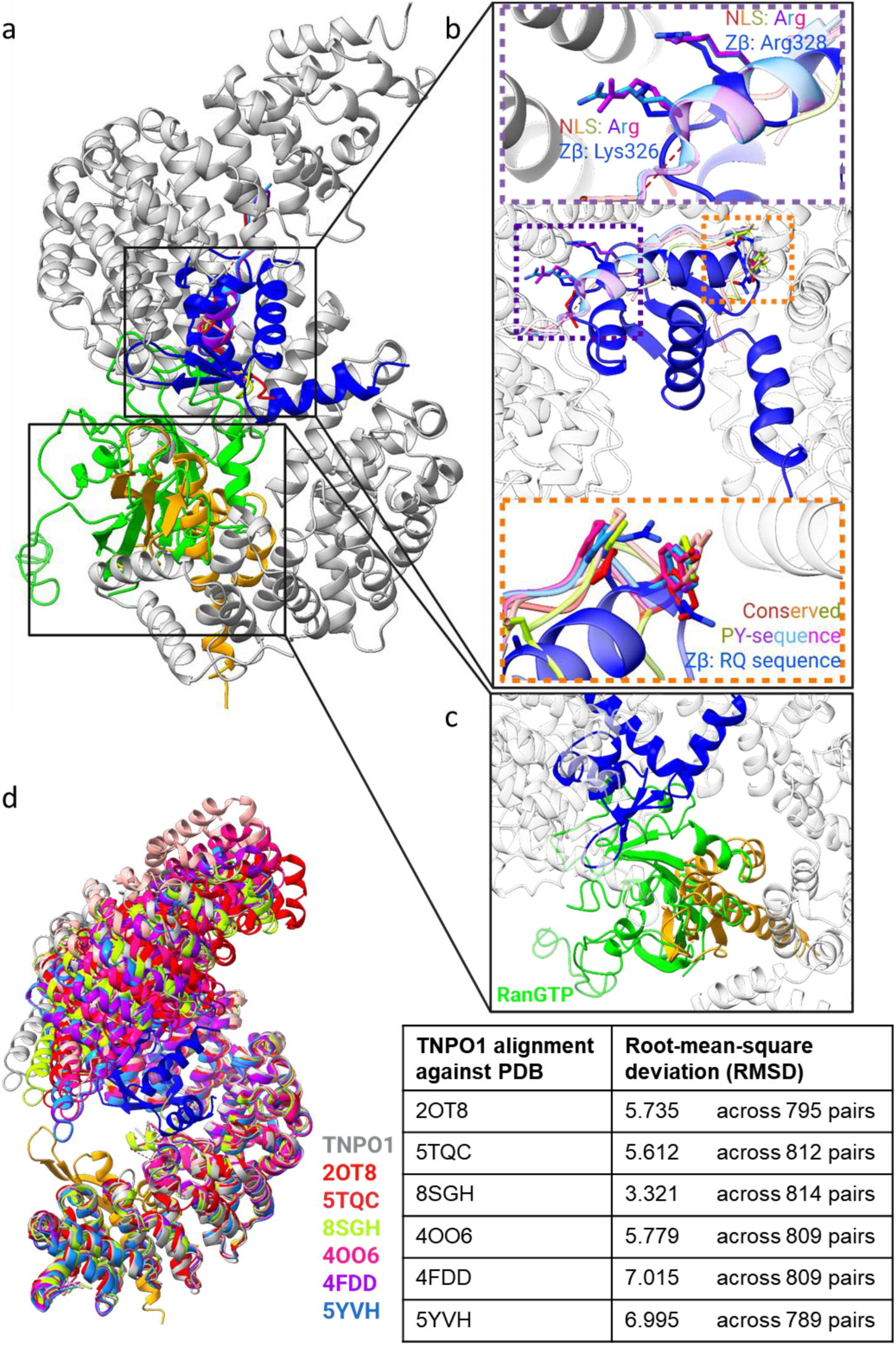
ADAR1 interacts with TNPO1 at both the canonical and non-canonical PY-NLS binding site. **a.** Alignment of ADAR1-TNPO1 complex with known cargoes of TNPO1. **b.** Zoomed in view of the alignment of Zβ domain and PY-NLS peptides (PDB 2OT8 in red, 5TQC in orange, 8SGH in lime green, 4OO6 in pink, 4FDD in purple and 5YVH in light blue) at the PY-NLS binding site. **c.** Zoomed in view of the alignment of dsRBD3 domain and Ran-GTP (PDB 1QBK in green). **d.** Alignment of TNPO1 against other TNPO1-PY-NLS complexes. The RMSDs are reported in the accompanying table. TNPO1 colored in grey, ADAR1 Zβ domain and dsRBD3 domain colored in blue and yellow respectively.

**Figure 4.**
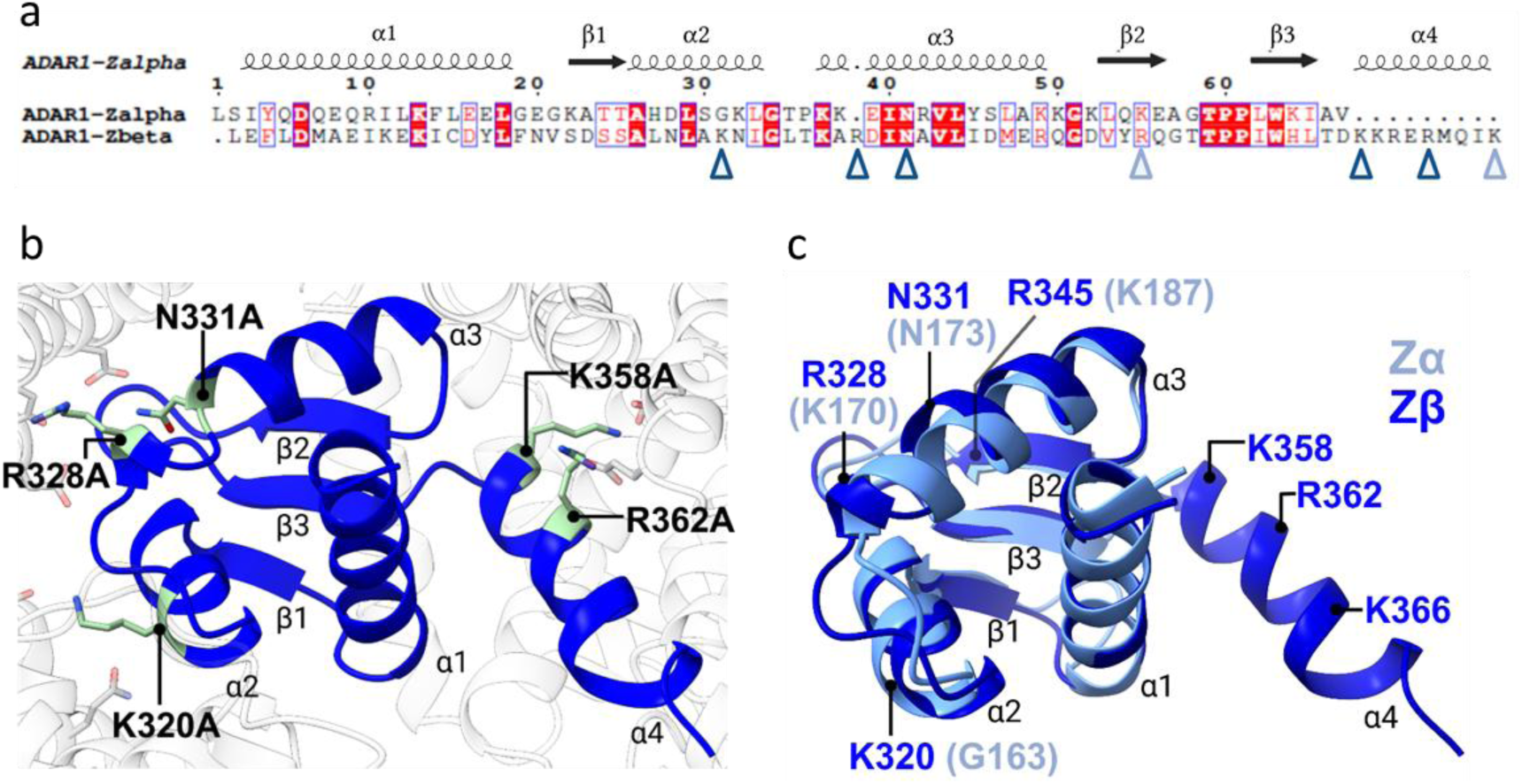
Comparison of ADAR1 Zβ and Zα domains. **a.** Multiple sequence alignment of ADAR1 Zα (L133-V199) and Zβ (F293-K366) binding domains. Arrows represent the critical interaction residues for TNPO1 recognition. Residues marked with a dark blue arrow were chosen for mutagenesis. **b.** 3D structure of ADAR1 Zβ with the mutation sites highlighted in teal. **c.** 3D structure alignment of ADAR1 Zβ with Zα (PDB 1QBJ). Critical interaction residues of ADAR1 Zβ are labelled in dark blue, with respective residues of ADAR1 Zα labelled in parentheses in light blue. ADAR1 Zα domain is modelled from L133 to V199, corresponding to the Zβ (F293-K366) domain. ADAR1 Zα and Zβ are coloured in light blue and dark blue, respectively.

Despite the absence of the characteristic PY dipeptide, the high density of arginine and lysine residues at the interface imparts an overall basic character consistent with PY-NLS features. Unlike the intrinsically disordered nature of most classical PY-NLS sequences, Zβ is a rigid and globular fold that is capable of occupying the same binding site through shape-mimicry and a charge-compatible surface. Notably, when TNPO1 was compared in its Zβ-bound and PY-NLS–bound states, a widening of the central groove was observed in the Zβ-bound complex (Fig. 3d, S4c), consistent with the conformational adaptability of TNPO1 in accommodating cargoes of different sizes and structures. These observations support a model in which Zβ functions as a structured, non-canonical TNPO1 recognition element, using its compact tertiary structure to emulate key recognition elements of linear PY-NLS motifs. This suggests that TNPO1 can accommodate not only flexible, linear signals but also structured domains that project compatible physicochemical properties. The conservation of Zβ across species and ADAR1 isoforms (Fig. S5) further supports its functional importance. Its strategic positioning within TNPO1’s central cargo cleft suggests that Zβ provides an additional anchoring interface during ADAR1 recognition by TNPO1.

Beyond this canonical interface, our structural model revealed that ADAR1’s third dsRBD engages TNPO1 at a secondary, non-canonical site located on the outer surface of the importin’s solenoid (Fig. 1d, Fig. 2b, 2d). This interface lies distant from the central groove and overlaps with the RanGTP-binding region observed in the TNPO1–RanGTP complex (PDB: 1QBK [35]), particularly across HEAT repeats 7 to 10 (Fig. 3a, 3c). The dsRBD3–TNPO1 interface buries approximately 1,106 Å² of solvent-accessible surface area (Table S2), representing an 18.4 % smaller interface relative to Zβ–TNPO1. This interface is characterized by a network of hydrogen bonds and van der Waals contacts involving residues distributed across the well-folded dsRBD3 domain (Fig. 2d). Additional contacts involving R736, D786, R790, V791, G794 and E795 further contribute to the interface with TNPO1 (Fig. 2d). Furthermore, residues E797 and K798 from the C-terminal flanking sequence contribute supplementary contacts with TNPO1, consistent with the previously identified role of this region in nuclear import. This spatial convergence suggests that dsRBD3 interaction and RanGTP binding may be functionally coupled, with RanGTP potentially displacing dsRBD3 during the nuclear import cycle [36]. Notably, this site is distinct from the classical PY-NLS binding cleft, reinforcing a model in which ADAR1-p110 employs a dual-site strategy to engage TNPO1.

### Mutations at the Zβ–TNPO1 interface abolish binding

To investigate the functional significance of the Zβ–TNPO1 interface, we performed targeted mutagenesis of residues within the Zβ domain identified by structural analysis as mediating TNPO1 binding. Five key residues—K320, R328, N331, K358, and R362 spanning helices 2, 3, and 4 of Zβ, were mutated in combination to generate four double mutants (Fig. 4a-b). The resulting constructs were designated as follows: ADAR1 Zβ WT (wild type), ADAR1 Zβ K320A/R328A, K320A/N331A, K320A/K358A, and K320A/R362A. These variants were tested for their ability to interact with full-length TNPO1 using biolayer interferometry (BLI). In these assays, biotinylated TNPO1 was immobilized onto Streptavidin-XT sensor probes, and binding to ADAR1-Zβ constructs was measured at concentrations up to 20 μM. Sensorgrams were recorded during both association and dissociation phases, and binding responses were analyzed using a 1:1 binding model.

Wild-type Zβ exhibited detectable binding to TNPO1, with an apparent dissociation constant (KD) of 80.6 nM (Fig. S7c). In contrast, mutant constructs displayed reduced or undetectable binding responses. The K320A/R328A and K320A/R362A mutants exhibited no detectable binding signal (Fig. S7d, g), indicating a complete loss of interaction. By contrast, the K320A/N331A and K320A/K358A mutants retained partial binding, albeit with markedly reduced binding affinity, with apparent KD values of 727 nM and 188 nM, respectively, in the initial screening assay (Fig. S7e, f). These findings indicate that the Zβ–TNPO1 interface is distributed across several residues, with the effects of individual double-mutant combinations differing in magnitude. In particular, the results suggest that K320, together with either R328 or R362, contributes substantially to the interaction. Based on these results, the double mutants, K320A/R328A and K320A/R362A, were selected for downstream functional assays.

Interestingly, despite their structural similarity, ADAR1 Zα also displayed detectable TNPO1 binding, but with a substantially weaker apparent KD of 635 nM, approximately eight-fold weaker compared to Zβ (Fig. S7b). Pairwise alignment and structural analysis suggest that this difference arises from the absence of several Zβ interface residues and the terminal α-helix (α4) present in Zβ, which provides additional residues at the TNPO1-binding interface (Fig. 4a, c). The disparity in affinity between these closely related domains suggests that TNPO1 recognition depends not only on the shared Z-domain fold but also on the specific composition and structural features of the binding surface. These distinctions highlight differences in the molecular recognition properties of the two homologous domains, with Zβ exhibiting stronger TNPO1 binding than Zα. This functional repurposing of the Zβ domain provides a molecular basis for its evolutionary conservation across isoforms and species, highlighting its pivotal role in maintaining ADAR1-p110 nuclear residency.

### Zβ mutations disrupt TNPO1-dependent nuclear import of ADAR1-p110 in cells

To assess the cellular consequences of disrupting the Zβ–TNPO1 interaction, the double mutations were introduced into full-length ADAR1-p110 isoform. In parallel, a Zβ deletion construct (ΔZβ) was generated to directly assess the contribution of the Zβ domain to ADAR1-p110 localization. The localization of wild-type and mutant ADAR1-p110 constructs was analyzed in HEK293T cells by transient transfection (Fig. 5). As expected, wild-type ADAR1-p110 localized predominantly to the nucleus. In contrast, deletion of the Zβ domain resulted in increased cytoplasmic localization. Similarly, both Zβ double mutants showed impaired nuclear localization and increased cytoplasmic accumulation, indicating that disruption of the Zβ domain compromises efficient nuclear localization of ADAR1-p110.

**Figure 5.**
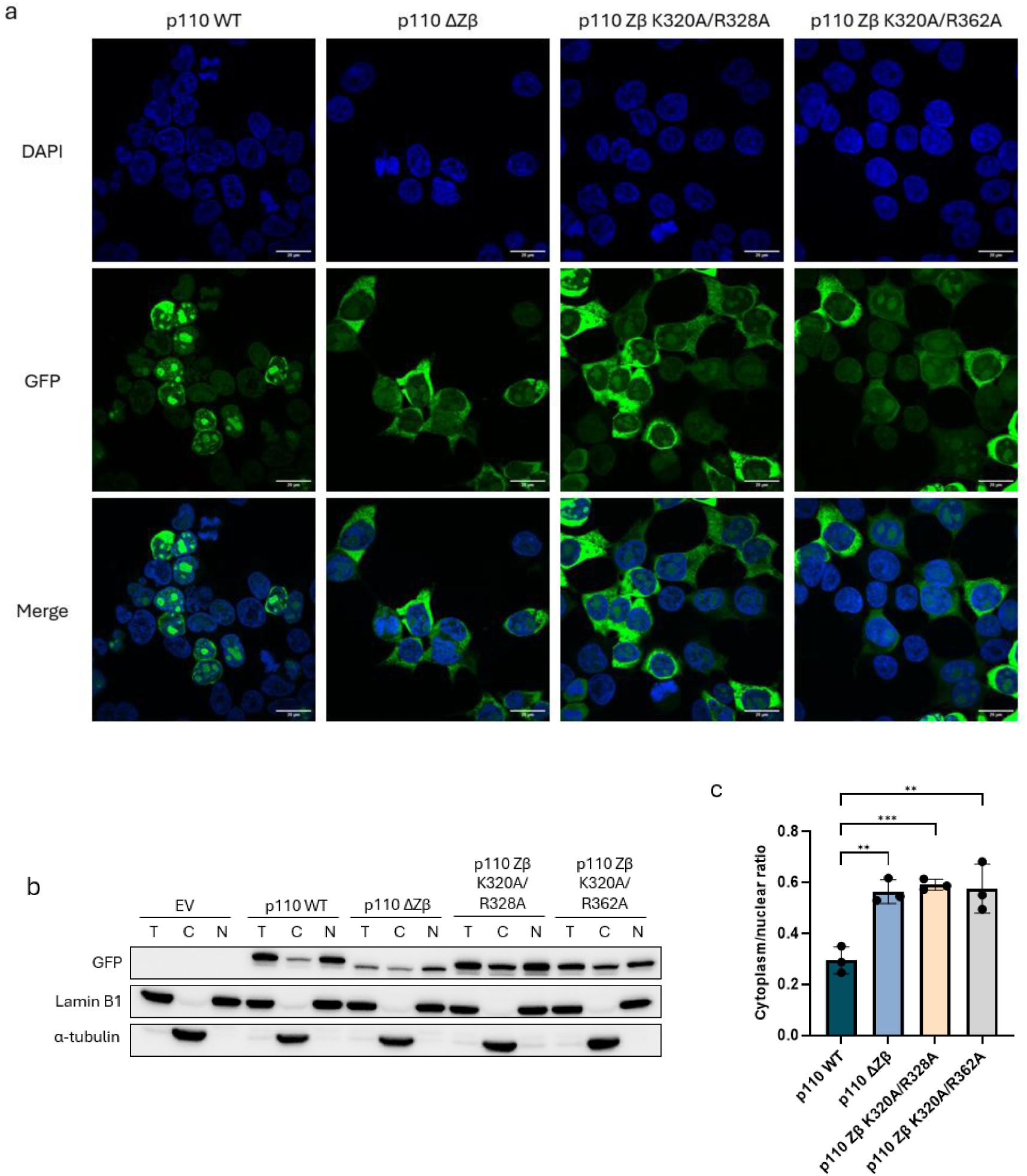
Mutation in the Zβ domain promotes cytoplasmic localization of ADAR1p110. **a.** Representative fluorescence images of HEK293T cells 24 hours post-transfection with GFP-tagged ADAR1p110 WT, ΔZβ, K320A/R328A, or K320A/R362A constructs. Nuclei were stained with DAPI. Scale bar represents 20 μm. **b–c.** Representative cell-fractionation western blot and quantification of the cytoplasmic-to-nuclear ratio of GFP-tagged ADAR1p110 WT and the indicated Zβ mutants from three independent experiments. Lamin B1 and α-tubulin were used as nuclear and cytoplasmic fraction markers, respectively. T, total lysate; C, cytoplasmic fraction; N, nuclear fraction. Data are shown as mean ± SD with individual biological replicates. Statistical significance is indicated as p < 0.05 (*), p < 0.01 (**), and p < 0.001 (***).

Consistent with the fluorescence imaging, cell fractionation showed approximately two-fold increase in cytoplasmic localization of the ΔZβ and Zβ mutant constructs compared with wild-type ADAR1-p110 (Fig. 5b–c). The two Zβ mutants showed broadly similar localization patterns, with only modest differences in the extent of cytoplasmic accumulation. Notably, these effects were observed despite the presence of the intact dsRBD3-flanking NLS, supporting an additional role for the Zβ domain in ADAR1-p110 nuclear import. As the Zβ point mutants recapitulated the cytoplasmic mislocalization observed with Zβ deletion, the mutant constructs were used for subsequent experiments to specifically investigate the functional consequences of disrupting the Zβ–TNPO1 interaction.

Together, these results support a collaborative mechanism in which the Zβ domain and the dsRBD3-associated NLS contribute to TNPO1-dependent nuclear import of ADAR1-p110. Disruption of the Zβ domain, either by deletion or point mutation, reduces efficient nuclear localization and promotes cytoplasmic accumulation of ADAR1-p110.

### ADAR1-p110 Zβ mutants enhance cytoplasmic immunosuppression

Given that ADAR1-p150 modulates innate immune responses, we asked whether altered localization of ADAR1-p110 caused by Zβ mutations affects its immunoregulatory function. HEK293T cells with stable ADAR1 knockout and MDA5 overexpression, expressing either wild-type or mutant forms of ADAR1 Zβ or p110 constructs were analyzed for activation of interferon-stimulated genes (ISGs) and cytosolic stress pathway markers. As expected, cells expressing mutant Zβ alone did not exhibit significant differences in the expression levels of cytosolic sensors such as PKR and IRF3 compared to wild type (Fig. 6). This is consistent with the understanding that Zβ alone is insufficient to modulate innate immune signaling. ADAR1 Zβ lacks the high-affinity dsRNA-binding capability and has not been shown to interact directly with innate immune receptors or accessory factors that mediate downstream signaling.

**Figure 6.**
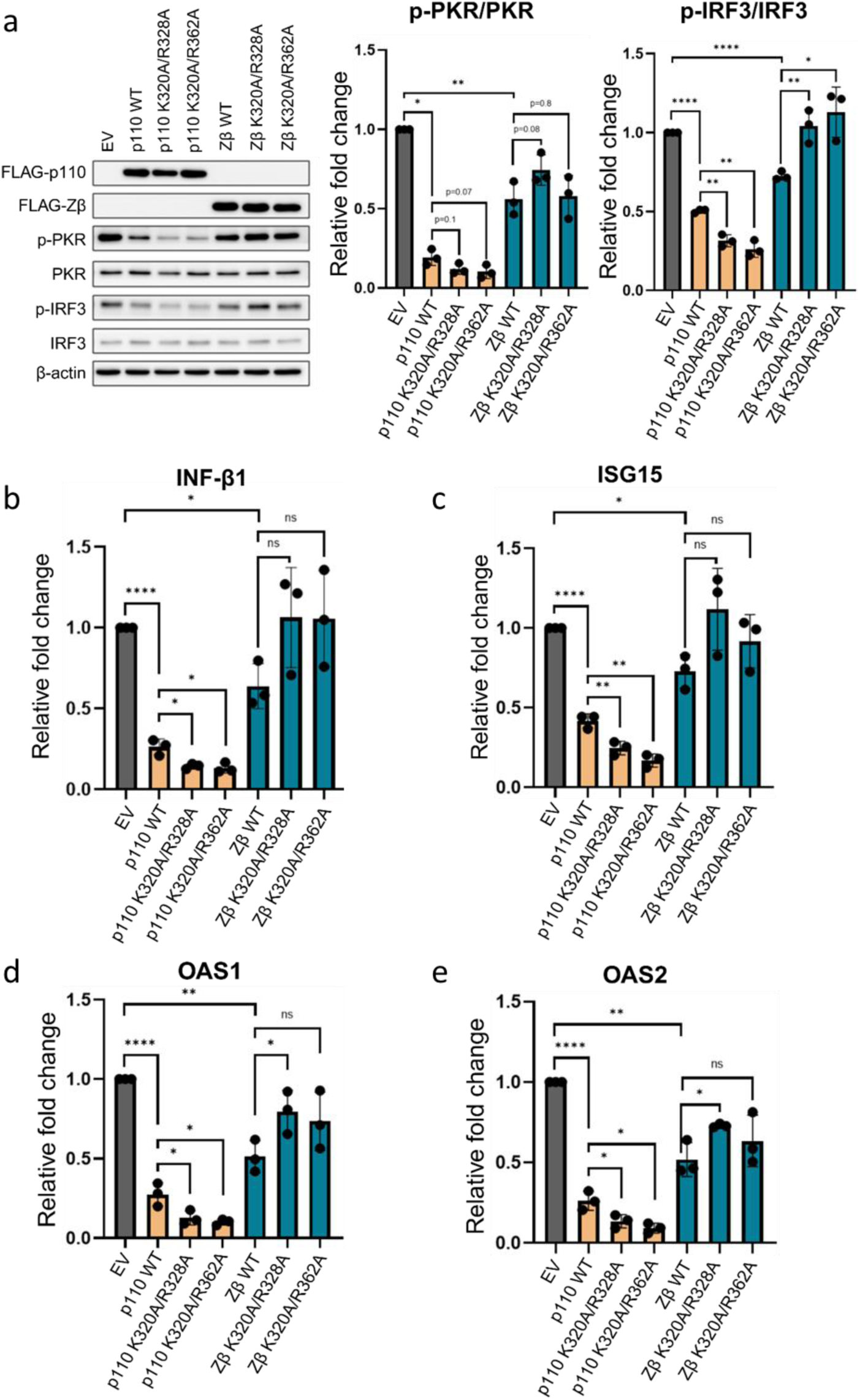
ADAR1-p110 mutants display increased immuno-suppression compared to the wildtype. **a.** Western blot analysis of overexpressed ADAR1p110 or Zβ wild-type or mutants on PKR and IRF3 phosphorylation. Relative fold change of phosphorylated-PKR/PKR and phosphorylated-IRF3/IRF3 reflected in the respective panels. Relative fold change of expression for **b.** Interferon-β1, **c.** ISG15, **d.** OAS1 and **e.** OAS2, analyzed by quantitative PCR. Statistical significance is indicated as p < 0.05 (*), p < 0.01 (**), p < 0.001 (***), and p < 0.0001 (****).

In contrast, cells expressing full-length ADAR1-p110 mutants showed a marked reduction in PKR and IRF3 activation (Fig. 6a). ISG expression—including OAS1, OAS2, and ISG15—was also diminished in the presence of the cytoplasm-localized p110 mutants compared to wild-type (Fig. 6b-e). This enhanced immunosuppressive activity correlated with the increased cytoplasmic retention of the mutants, suggesting that altered subcellular localization may influence the immunoregulatory activity of ADAR1-p110. Although previous studies have implicated ADAR1-p110 in the regulation of innate immune signaling through editing-independent mechanisms, including protein–protein interactions involving the dsRBDs [4, 37], the contribution of RNA-editing activity to the observed phenotype remains unclear. As RNA-editing activity was not assessed in this study, it remains possible that the enhanced suppression of innate immune signaling arises from altered RNA-editing activity, editing-independent interactions or a combination of these mechanisms.

Interestingly, even mutants exhibiting only modest defects in nuclear import could elicit strong ISG suppression, raising the possibility that even relatively small changes in cytoplasmic ADAR1 abundance could influence innate immune signaling. These findings suggest a functional relationship between the structural interaction of Zβ with TNPO1 and the immunoregulatory role of ADAR1-p110. Altogether, our data emphasize the importance of spatial regulation in modulating ADAR1 function: while nuclear localization enables RNA editing, cytoplasmic accumulation— induced through disruption of the Zβ–TNPO1 interface—repositions ADAR1-p110 as an effective suppressor of innate immune activation. This dual functional capacity reinforces the critical role of Zβ in regulating both the localization and immune activity of ADAR1-p110, with direct implications for understanding how its dysregulation may contribute to immune-related pathologies.

## Discussion

Our study reveals a previously unrecognized mechanism by which the Zβ domain of ADAR1-p110 contributes alongside dsRBD3 to direct subcellular localization and influence immune regulatory function via the nuclear import receptor TNPO1. Through high-resolution cryo-EM analysis, we demonstrate that Zβ and dsRBD3 form a composite TNPO1-binding interface involving two spatially distinct regions of the import receptor. The Zβ domain anchors into TNPO1’s central cargo-binding groove via a positively charged surface that partially mimics features of classical PY-NLSs, while dsRBD3 engages a non-canonical outer surface overlapping the RanGTP binding site. This dual-domain interface underscores the adaptability of TNPO1 in recognizing both structured and disordered cargo-associated recognition elements.

An NMR-based molecular model of TNPO1 interaction was proposed by Barraud et al. in 2014 [17], who identified a bipartite PY-NLS comprising a disordered string of residues M708–K715 and E797–R801 flanking the dsRBD3 domain. They demonstrated that these flanking regions alone were sufficient to mediate nuclear import through the conventional cargo-binding groove. In contrast, our structure shows that dsRBD3 binds to TNPO1 as a folded domain at a peripheral site near the N-terminus, with the structured domain contributing directly to the peripheral interaction surface (Fig. 1d, 2b, 2d). The interface is stabilized by residues within the structured core of dsRBD3 as well as residues from the C-terminal flanking sequence that contribute supplementary contacts for enhanced stability of the interaction. These findings suggest that the structured dsRBD3 domain may contribute more directly to TNPO1 recognition than previously appreciated. Our findings and those of Barraud et al., therefore, highlight alternative modes of TNPO1 recognition. While the classical PY-NLS identified by Barraud may represent one mechanism of TNPO1 engagement, particularly with isolated constructs, our structure illustrates a complementary model in which ADAR1-p110 engages TNPO1 through a composite, structured-domain-mediated interface. This includes Zβ anchoring within the central cavity and dsRBD3 associating at a distinct, RanGTP-sensitive peripheral site. This bipartite interaction provides a structural framework for understanding how multiple regions of ADAR1 may contribute to TNPO1 recognition and nuclear import.

Structure-guided mutagenesis of Zβ residues that contact TNPO1 resulted in cytoplasmic mislocalization of ADAR1-p110, even in the presence of an intact dsRBD3 NLS. This altered localization was accompanied by changes in innate immune signaling readouts. While Zβ mutants alone had no detectable effect on cytosolic stress markers—consistent with their lack of dsRNA-binding or catalytic function—ADAR1-p110 mutants exhibited a marked suppression of PKR and IRF3 activation, and decreased expression of interferon-stimulated genes (ISGs) such as OAS1, OAS2, and ISG15. These findings suggest that cytoplasmic ADAR1-p110 retains functional capacity to suppress innate immune signaling, either by editing of immunogenic dsRNA, through editing-independent mechanisms, or a combination of both. Future investigations using catalytically inactive ADAR1-p110 mutants could help distinguish between these possibilities.

This observation contrasts with previous assumptions that nuclear exclusion of ADAR1 primarily drives immune activation, as seen in certain Aicardi–Goutières syndrome (AGS) mutations and ADAR1 knockout models [38, 39]. Our findings suggest that the consequences of altered ADAR1 localization may depend on isoform, cellular context and experimental conditions. In particular, the increased cytoplasmic accumulation of ADAR1-p110 was associated with suppression of innate immune signaling in our experimental system. In line with this finding, a study performed using mouse fibroblast cells deficient in ADAR1 suggested that ectopic overexpression of ADAR1-p110 is capable of suppressing ISGs similar to ADAR1 p150 [40]. In a separate study, an ADAR2 construct carrying a mutation that abolished nuclear localization also showed immunosuppressive effects, further substantiating the link between subcellular localization and immune signaling functions [41]. This effect could be further explained by studies showing that ADAR1-p110 can suppress PKR activation via direct interactions between its dsRBD3 and PKR’s kinase domain [4]. Furthermore, ADAR1 has been shown to promote Enterovirus D68 replication through its deaminase domain and inhibition of the PKR pathway [37], highlighting the dual role of ADAR1-p110 as both editor and immune suppressor. The Zβ–TNPO1 interface therefore may contribute to the spatial regulation of ADAR1-p110, influencing the distribution of ADAR1 between the nuclear and cytoplasmic compartments.

Comparative structural analysis with other Z-DNA binding domains (ZBDs), including ADAR1 Zα and the ZBP1 ZBDs, further underscores the specificity of this interaction between ADAR1 Zβ and TNPO1. Sequence comparisons (Fig. 4a, S6a) reveal that these related domains lack several residues identified at the Zβ-TNPO1 binding interface. Structural comparisons of ADAR1 Zβ with ADAR1 Zα (Fig. 4c) and ZBP1 ZBDs (Fig. S6b-c) further demonstrate that only ADAR1 Zβ possesses a terminal α-helix that contributes three key interacting residues, which may contribute to the observed surface complementarity with TNPO1. These observations suggest that among the two human ZBD-containing proteins, ADAR1 and ZBP1, ADAR1 Zβ has evolved distinct structural adaptations that enable functional recognition by TNPO1, reflecting its specialization in regulating ADAR1-p110 localization.

Furthermore, no disease-associated natural mutations have been reported within the Zβ domain, although the absence of reported mutations does not establish an essential role in nuclear localization. The immune suppressive effects observed are therefore best understood in the context of altered ADAR1-p110 subcellular distribution, although precise mechanism underlying these effects remains to be determined. Importantly, ADAR1 localization is dynamically regulated under physiological conditions. Under cellular stress, ADAR1-p110 is phosphorylated and exported to the cytoplasm to block apoptosis [16, 42]; During viral infection, both isoforms can accumulate in the cytoplasm to facilitate viral replication [37, 43]. These contexts highlight how dynamic regulation of ADAR1 localization becomes critical under perturbation.

The broader significance of Zβ-mediated localization is underscored by cancer studies showing that loss of ADAR1 induces viral mimicry programs and innate immune activation, with combined DHX9 and ADAR1 deficiency enhancing dsRNA-sensing–driven cell death in breast cancer cells [44]. Modulating ADAR1 localization via the Zβ domain could therefore influence immune surveillance and therapeutic responses. The Zβ–TNPO1 axis represents a potential target for tuning ADAR1 activity: whether modulation of this interaction could influence ADAR1-dependent immune regulation in disease contexts remains an interesting possibility for future investigation. Context-dependent regulation may also involve post-translational modifications, competitive TNPO1 interactions, or stress signaling. Collectively, our findings identify Zβ as an additional determinant of ADAR1-p110 subcellular localization through its interaction with TNPO1, establishing a structural framework for understanding the regulation of ADAR1 trafficking and its influence on innate immune signaling.

## Materials and Methods

### Plasmid generation

Plasmids for the bacterial ADAR1-ZBDRBD construct were purchased from Protein Production Platform, NTU. A secondary TwinStrep tag was cloned to the C-terminal of the ADAR1-ZBDRBD construct via InFusion cloning following manufacturer’s protocol. Wild-type ADAR1-Zβ was cloned into the same vector via end-to-end ligation. Mammalian expression plasmid for ADAR1 p110 and wild-type Zβ were obtained from Dr Polly Chen’s lab. Wild-type ADAR1-Zβ mutant constructs were generated sequentially (K320A followed by the second mutation) by site directed mutagenesis. Primer sets for preparing the constructs are listed in Table 1. Bold indicates the sites that are mutated.

**Table 1.**
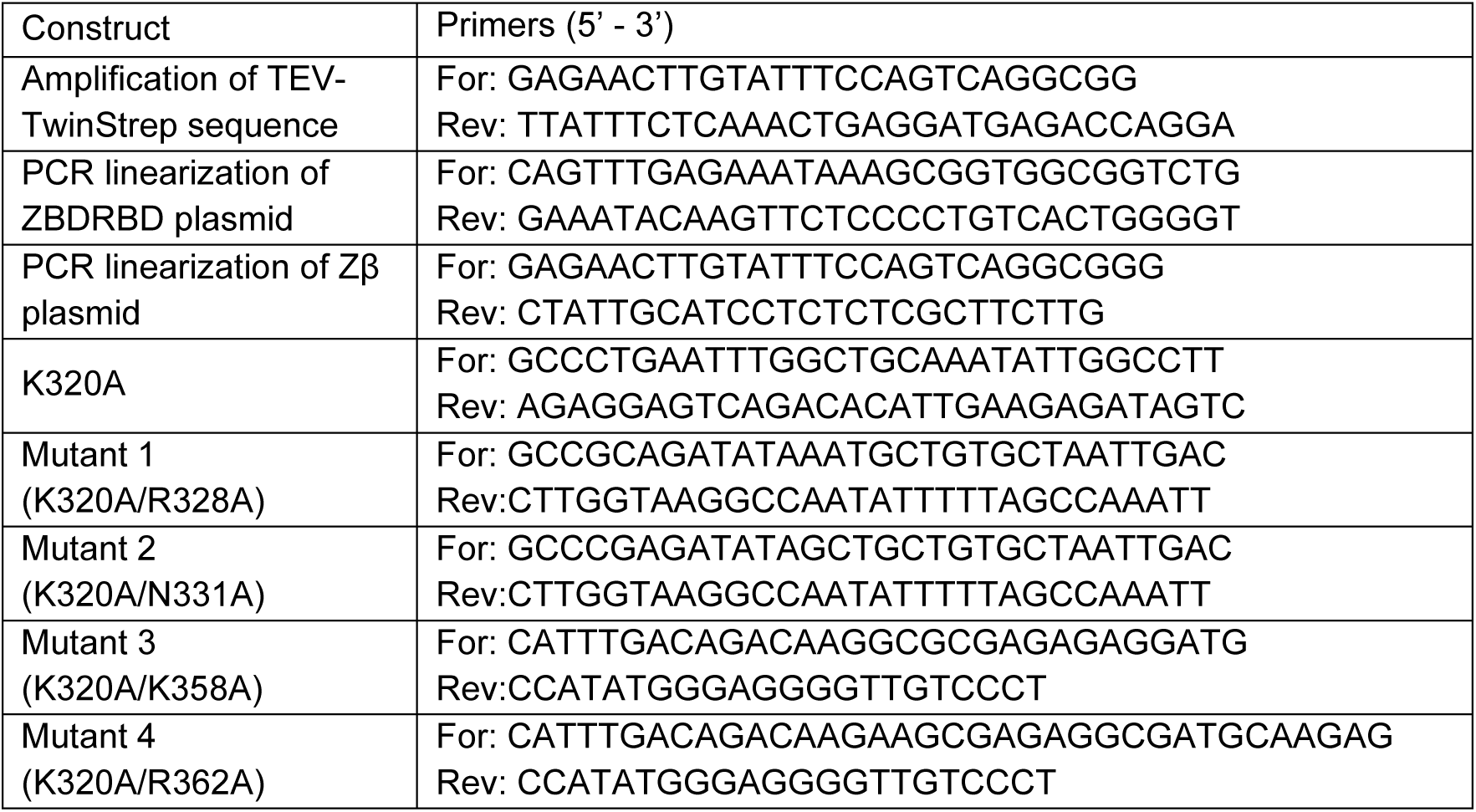
Forward and reverse primers for amplification of target sequence for cloning.

### Protein Expression

Plasmids were transformed into BL21 (DE3) Rosetta T1R E. coli strain. Protein expression was performed in LB Broth, Miller with appropriate antibiotics. Bacteria cells were grown at 37 ⁰C until OD600 of approximately 0.4, cooled to 18 ⁰C before induction with 1mM Isopropyl-β-D-thiogalactopyranoside (IPTG) at OD600 of approximately 0.8 to 1.0. Cells were harvested 18 to 20 hours post induction.

### Protein Purification

In brief, bacteria cell pellets are resuspended in lysis buffer containing 50 mM Tris-Cl pH 8.0, 300 mM NaCl, 0.1 % Triton-X100, 1 mg/mL lysozyme, 10 % glycerol, 10 mM β-mercaptoethanol, benzonase nuclease, RNAseA and 1 mM PMSF, and incubated on a roller platform for 1 hour before further lysis by sonication (40 % amplitude, 5 seconds on/ 10 seconds off, 4 ⁰C) for 5 minutes twice. The lysate was clarified by centrifugation. Clarified supernatant was incubated for one hour with Buffer A (25 mM Tris-Cl pH 8.0, 5 % glycerol, 10 mM β-mercaptoethanol) pre-equilibrated Ni-NTA or GST beads at 4 ⁰C with gentle rolling and loaded onto Econo-Pac Gravity Column. Beads were washed in five column volumes (CV) each of high salt wash buffer 1 (Buffer A with 1 M NaCl), followed by wash buffer 2 (Buffer A with 150 mM NaCl). Proteins were eluted with elution buffer (Buffer A with 500 mM imidazole or 10 mM gluthathione reduced) in five to 10 fractions of one CV each, with 10 minutes incubation between each elution. Eluted protein was cleaved overnight with the appropriate protease. Reverse pulldown was performed to separate cleaved and uncleaved protein products and concentrated to a suitable volume for loading onto HiTrap Heparin HP (GE Healthcare). Protein was eluted with a salt gradient over 10 column volumes in a buffer containing 20 mM HEPES pH 7.2, 10 % glycerol, 2 mM TCEP and either no or 1 M NaCl. Fractions containing the protein of interest were concentrated and then purified using HiLoad Superdex200 10/300 pg in SEC buffer containing 20 mM HEPES pH 7.5, 150 mM NaCl, 2 mM TCEP and 5 % glycerol. Purified proteins were collected and concentrated for downstream structural analysis. Purification was done on an ÄKTA Pure purification system (GE Healthcare) unless otherwise stated.

### Cryo-EM sample and grid preparation

Purified recombinant ADAR1-ZBDRBD protein was incubated with purified full length TNPO1 at a 1:1 ratio in SEC buffer for an hour on ice. The resulting mixture was passed through analytical size exclusion chromatography using Superdex 200 3.2/300 Increase column. Peak fractions containing both proteins were collected for cryo-grid preparation.

Cryo-grid preparation was performed with the Vitrobot Mark IV (Thermo Fisher Scientific). Cryo-grids used were 300-mesh gold grids with R1.2/1.3 holey carbon film (Quantifoil, EMS). The grids were glow discharged for 1 minute in air plasma immediately before use. 2.5 µL sample was pipetted onto the grid surface, blotted for 4 seconds and immediately plunge frozen in liquid ethane. The grids were subsequently clipped for data collection.

### Cryo-EM data collection and processing

The dataset of TNPO1-ADAR1 complex was collected in-house (FACTS, NTU). 5,000 micrographs were recorded on the Titan Krios electron microscope operated at 300 kV. Detailed parameters for the data collection are summarized in Supplementary Table S1. The micrographs were motion-corrected on CryoSPARC Live. The motion corrected micrographs were then imported into CryoSPARC-v4.5 for further processing. Using a template of TNPO1 (EMD 40455), 9,064,647 particles were picked and extracted. The particles were subjected to three cycles of 2D classification to obtain 71,006 particles that were used for Topaz training with the settings (10 epochs and 5000/epoch). 1,623,957 particles were re-extracted. 2D classification yielded 1,084,497 selected particles, separated into five classes by ab-initial reconstruction. These five classes were then subjected to heterogenous refinement, with one class showing clear protein density selected for further processing. The particles from the good class were subjected to multiple iterations of 3D classification, non-uniform and local refinement, and CTF refinements. A final 3.15 Å map with 56,431 particles was obtained after a final local refinement.

The model of full length TNPO1 was downloaded from Protein Data Bank (PDB 2QMR). For ADAR1, the model of the dsRBD3 was downloaded from PDB 2MDR. The model for the remainder unknown density was rebuilt using ModelAngelo supplemented with the FASTA sequence of ADAR1-ZBDRBD. Each model was docked into the density map in ChimeraX and manually refined using the ISOLDE plugin. The model was further refined in Phenix with real-space refinement and secondary structure and geometry restraints. All figures were generated by UCSF ChimeraX.

### Structural Interface Analysis

Buried solvent-accessible surface areas (BSAs) at the protein–protein interfaces were calculated using UCSF ChimeraX with the “measure buriedarea” command. The interacting protein chains were specified individually using the syntax “measure buriedarea #x/y with #m/n”, where *x* and *m* denote model identifiers and *y* and *n* denote chain identifiers, respectively. The resulting interface areas were reported in Å^2^ in Supplementary Table S2.

### Biolayer Interferometry

Purified TNPO1 protein was biotinylated and desalted using the EZ-Link Sulfo-NHS-LC-Biotinylation kit (Thermo Fisher Scientific) at a molar ratio of 1 protein to 2 biotin following manufacturer protocols. Analyte proteins containing the TwinStrep tag was cleaved with TEV protease overnight at 4 ⁰C. Reverse pulldown was performed to obtain cleaved proteins for kinetics assay.

In brief, Streptavidin-XT (SA-XT) probes (Gator Bio) were hydrated in binding buffer (20 mM HEPES pH 7.5, 150 mM NaCl, 0.02 % Tween-20, 0.05 % BSA) for at least 10 minutes at room temperature prior to the start of the assay. Biotinylated TNPO1 was diluted to 0.2 µM and aliquoted to the sample plate. Analyte proteins were diluted to a 20 µM starting concentration (screening run) and further diluted two-fold to obtain a 5-concentration series (titration run). The kinetics assays were executed on the Gator Prime system (Gator Bio) with the following settings: 1000 rpm shaking at 30 ⁰C. The run is proceeded as follows: 1. First baseline: 120 seconds, 2. Loading: 180 seconds, 3. Second baseline: 120 seconds, 4. Association: 150 seconds, and 5. Dissociation: 150 seconds. All data analysis was performed on the Gator Navigator software (Gator Bio). The data were aligned to the beginning of the association phase with inter-step correction. All data points were subtracted against the reference biosensor loaded only with the ligand. A global 1:1 fitting model with the Rmax Unlinked was applied to the titration series.

### Cell culture

HEK293T cells were cultured in DMEM high glucose media (Biowest) supplemented with 10 % fetal bovine serum (FBS; Thermo Scientific) and incubated at 37 °C in a humidified incubator containing 5 % CO_2_.

### Cell imaging

HEK293T cells were transfected with 1 μg of GFP-tagged plasmid encoding either full-length ADAR1 p110 WT or ΔZβ, K320A/R328A, or K320A/R362A constructs, using Lipofectamine 2000 at a 1:3 DNA-to-reagent ratio. Cells were incubated for 24 hours post-transfection prior to staining with DAPI. Confocal imaging was performed at 100x magnification.

### Generation of ADAR1-KO MDA5-overexpressed cells

HEK293T cells were transfected with a Lenti-CRISPR vector encoding an ADAR1-targeting sgRNA (AATAGTATCCGCGCAGCACC) and subsequently subjected to single-cell isolation. Genomic DNA from individual clones was extracted, and the target locus was PCR-amplified and submitted for Sanger sequencing. Indels were analyzed using TIDE (https://apps.datacurators.nl/tide/), and candidate clones were further validated by western blot to confirm loss of ADAR1 protein expression. The clone with validated loss of ADAR1 protein were transduced with CSII-CMV-MCS-IRES2-Venus containing MDA5 sequence, followed by puromycin selection to generate ADAR1-KO MDA5-overexpressed cells.

### Western blot

Protein lysates were extracted using RIPA buffer (Sigma) supplemented with 1× cOmplete EDTA-free protease inhibitor cocktail (Roche) and 1x PhosSTOP™ (Roche). Lysates were resolved on 10 % SDS–PAGE gels, transferred onto PVDF membranes, and incubated with primary antibodies overnight at 4 °C, followed by incubation with secondary antibodies at room temperature for 1 hour. Primary antibodies used in this study are anti-FLAG M2 antibody (Cell Signalling Technology, 86861), anti p-PKR (Abcam, ab32036), anti-PKR (Santa Cruz Biotechnology, sc-6282), anti p-IRF3 (Abcam, ab76493), anti-IRF3 (Santa Cruz Biotechnology, sc-33641), anti-GAPDH (Santa Cruz Biotechnology, sc-59540).

### Cell fractionation

Nuclear and cytoplasmic fractions were isolated using the Nuclear Extract Kit (Active Motif, Carlsbad, CA) following the manufacturer’s instructions. Equal amount of nuclear and cytoplasmic fractions was used for western blot analysis. Fraction purity was assessed using anti–Lamin B (Proteintech, 12987-1-AP) for the nuclear fraction and anti–α-tubulin (Proteintech, HRP-66031) for the cytoplasmic fraction.

### Quantitative PCR

Total RNA was isolated using the RNeasy Mini Kit (Qiagen), followed by reverse transcription into complementary DNA (cDNA) using the SensiFAST™ cDNA Synthesis Kit (Bioline). Quantitative PCR (qPCR) was then carried out with GoTaq qPCR Master Mix (Promega). Primer sequences are listed in Table 2.

**Table 2.**
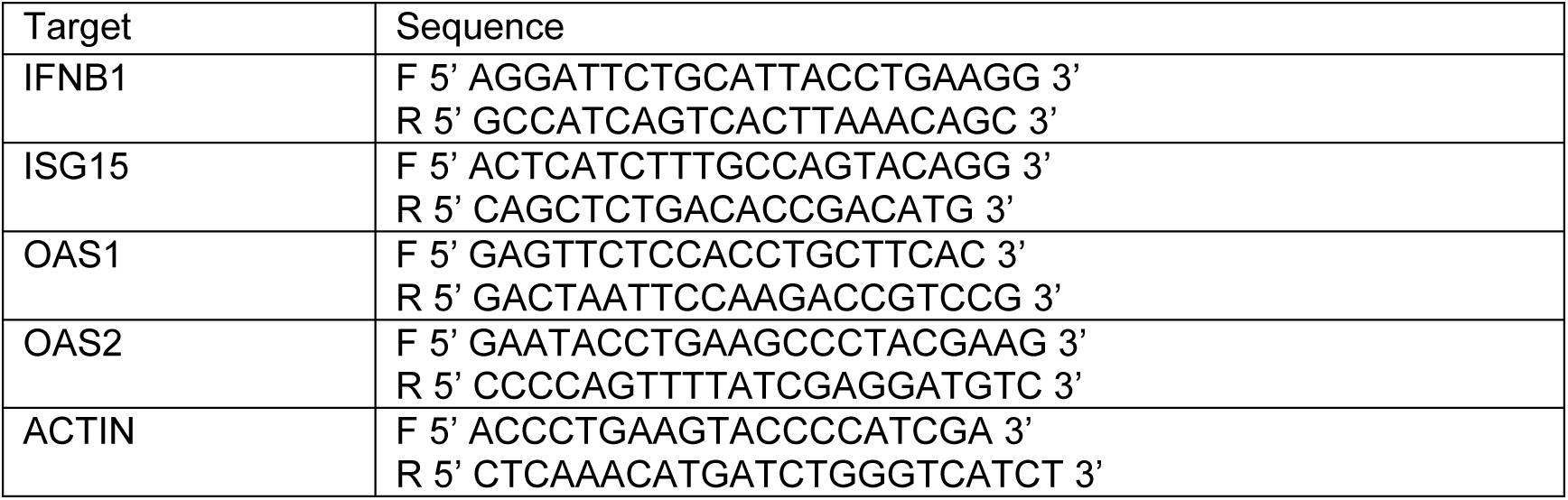
Primer sequences for quantitative polymerase chain reaction (qPCR) amplification of target genes.

### Multiple Sequence Alignment

The sequences of ADAR1 protein of various species were obtained from Uniprot website and aligned using ClustalOmega [45]. For alignment of the ADAR1 and ZBP1 Z-DNA binding domains, alignment was modified using ESPript [46] to obtain the structure-based alignment.

### Statistical Analysis

Western blot and qPCR data were analyzed using unpaired, two-tailed Student’s t-test. Data are presented as mean ± SD (n = 3 biological replicates). Statistical significance is indicated as p < 0.05 (*), p < 0.01 (**), p < 0.001 (***), and p < 0.0001 (****).

## Acknowledgments

This research is supported by the Singapore Ministry of Education under its Singapore Ministry of Education Academic Research Fund Tier 3 (MOET32023-0003) to PC and Education Academic Research Fund Tier 1 (RT22/23) to DL. We thank Dr. Lu Lei for providing the TNPO1 expression plasmid. We thank the scientific facility support from NTU Institute of Structural Biology and Protein Product Platform. We thank members of the DL and PC lab for their support.

## Data And Code Availability

The SPA-cryoEM density map of the ADAR1-TNPO1 complex has been deposited in EM Database under the accession code EMD-100045. The corresponding atomic coordinates have been deposited in the Protein Data Bank under accession code 34NN.

