## Supplementary Information for "Mechanistic Insights into ADAR1-p110 Nuclear Import: A Dual-Interface Interaction with Transportin-1"

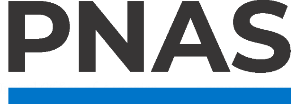

**Supporting Information for**

Mechanistic Insights into ADAR1-p110 Nuclear Import: A Dual-Interface Interaction with Transportin-1

Danielle Huey Ren Chin ^a^, Jesslyn Zhou ^c^, Yongbo Luo ^a^, Davin Hong ^a^, Yaw Bia Tan ^a #^, Polly Leilei Chen ^c d e^ *, Dahai Luo ^a b f^ *

* To whom correspondence may be addressed: Associate Professor Luo Dahai or Associate Professor Polly Chen. **Email:**

**This PDF file includes:**

Supplementary Figures S1-S7

Supplementary Table S1-S2

Figures

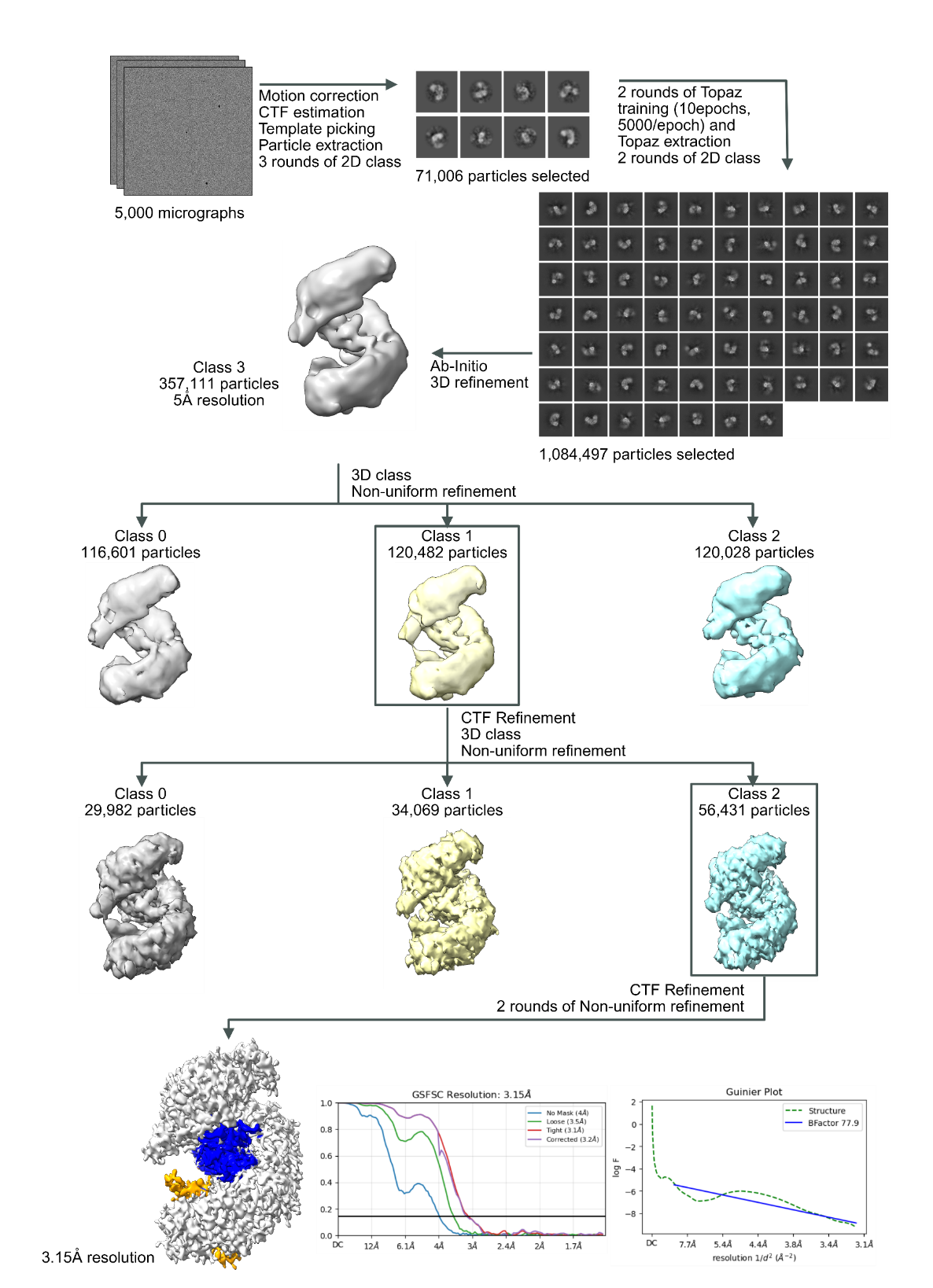
Fig. S1. Cryo-EM data processing workflow for the reconstruction of the complex between TNPO1 and ADAR1. The pipeline shows representative micrographs collected at 165,000x magnification, 2D classes, 3D reconstructions and refinement steps. Validation of cryo-EM structure of the complex between TNPO1 and ADAR1 are shown using the Gold-standard Fourier shell correlation (GSFSC) curves and Guinier Plot for the final map. Masked (tight mask in red, loose mask in green), unmasked (blue), and corrected mask (purple) plots are shown. All data processing was completed using CryoSPARC-v4.5.

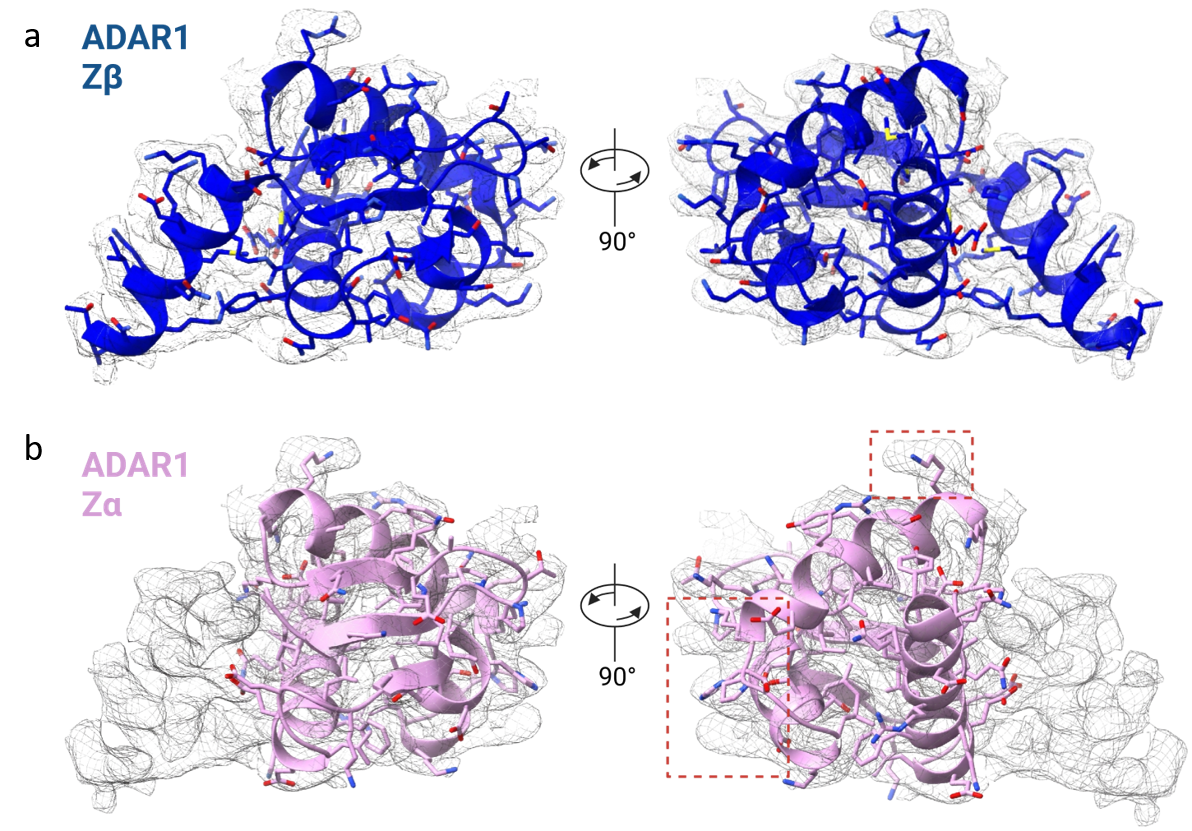
Fig. S2. Local domain comparison of a. ADAR1 Zβ and b. Zα domain, within the density map. The red dashed boxes indicate regions where the residues do not correspond to the density map. ADAR1 Zβ domain and Zα domain coloured in blue and pink respectively.

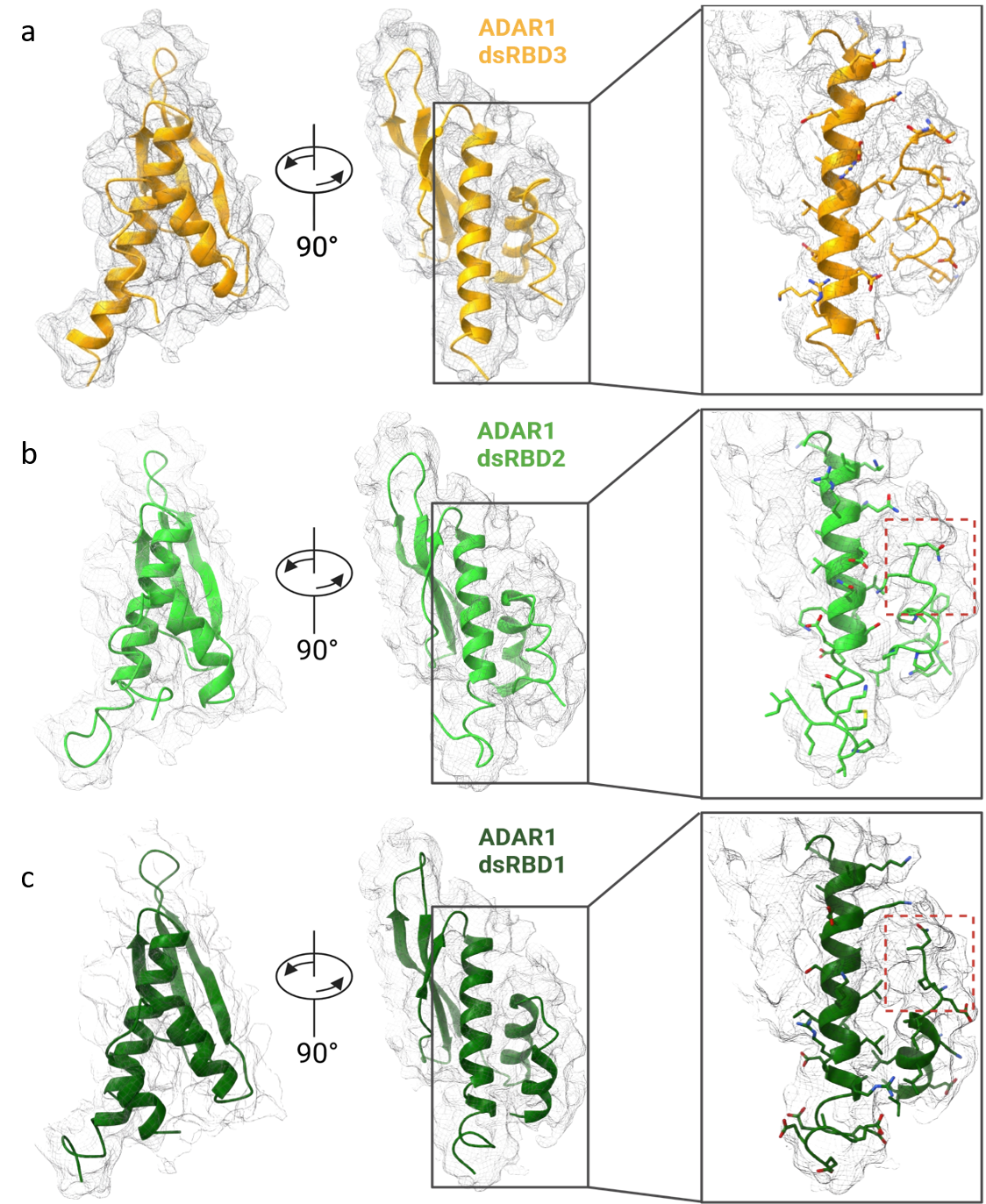
**Figure S3.** Local domain comparison of **a**. ADAR1 dsRBD3, **b.** ADAR1 dsRBD2, and **c.** ADAR1 dsRBD1 domains within the density map. The region corresponding to the nuclear localization sequence in dsRBD3 is highlighted in the grey box. The red dashed boxes indicate regions where the residues do not correspond to the density map. ADAR1 dsRBD3 domain coloured in yellow, ADAR1 dsRBD2 in lime green, and ADAR1 dsRBD1 in forest green respectively.

**Figure S4.** Structure model alignment of **a.** ADAR1 Zβ domain, **b.** ADAR1 dsRBD3 domain, and **c.** TNPO1, against their respective crystallographic or NMR models. TNPO1 colored in grey, ADAR1 Zβ domain and dsRBD3 domain colored in blue and yellow respectively.

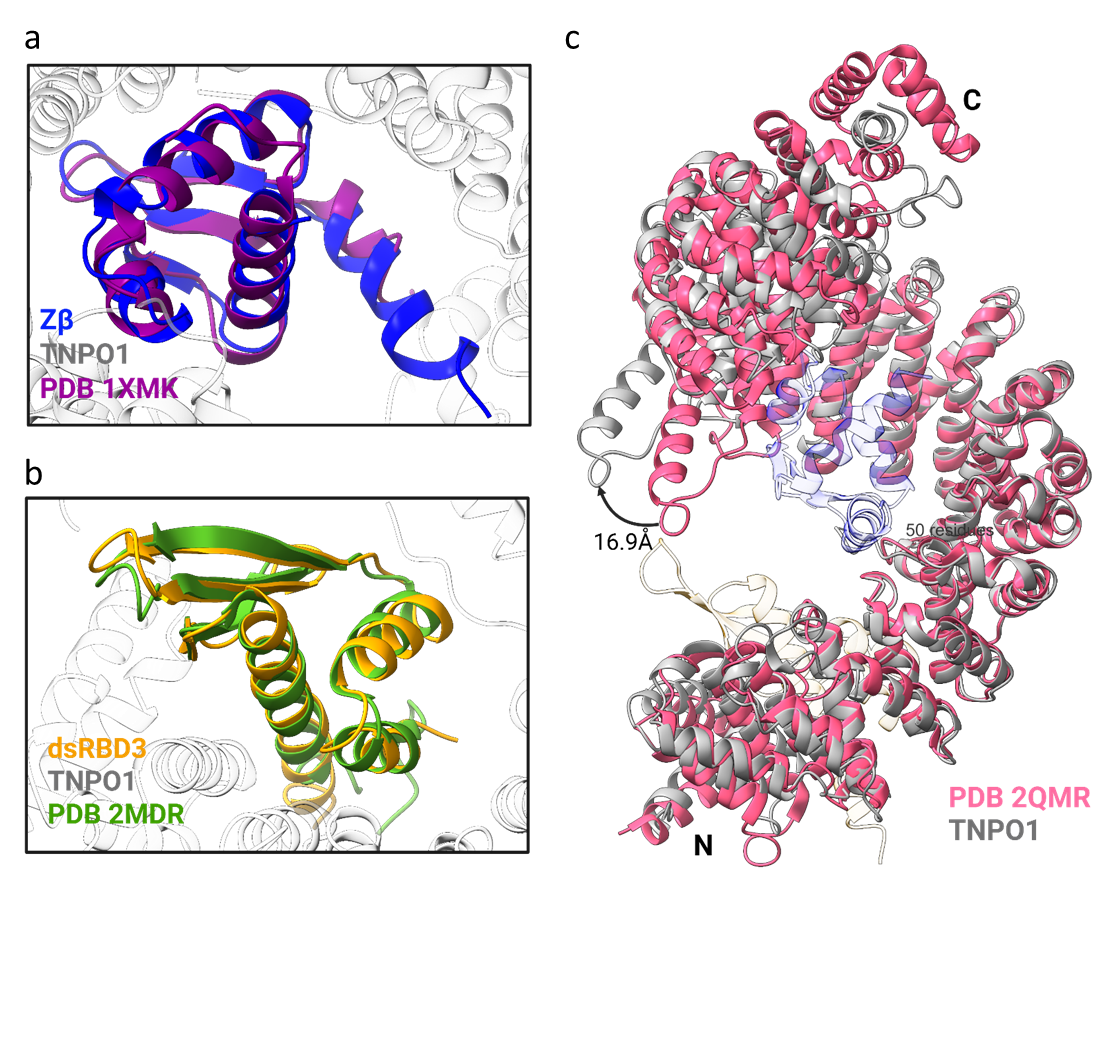

**Figure S5.** Multiple sequence alignment (MSA) of ADAR1 across multiple species. Region shown encompasses the Zβ domain (F293 – K366). Alignment is annotated based on the numbering of human ADAR1-p150. Arrows represent the interaction residues chosen for mutagenesis. MSA prepared using ClustalOmega.

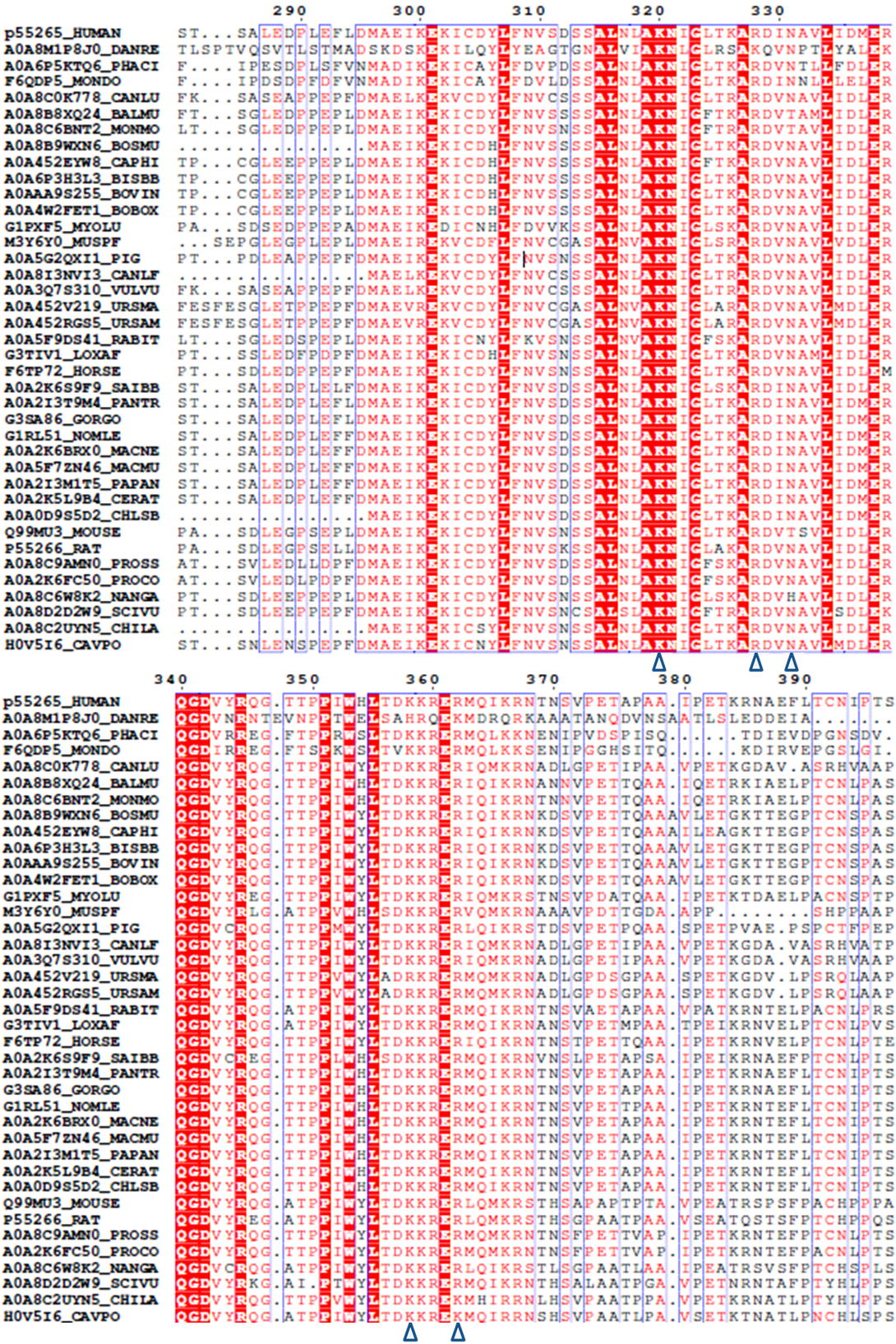

**Figure S6.** Alignment of ADAR1 Zβ domain against ZBP1 Z-DNA binding domains. **a.** Multiple sequence alignment of ADAR1 Zβ domain to ZBP1 Zα1 (P80-G70) and Zα2 (P103-R166). Arrows represent the critical interaction residues between ADAR1 Zβ and TNPO1. 3D structure alignment of **b.** ADAR1 Zβ to ZBP1 Zα1 (PDB 9J89), and **c.** ADAR1 Zβ to ZBP1 Zα2 (PDB 3EYI) are shown. ZBP1 Zα1 and Zα2 are modelled from P9 to G70, and P103 to R166, respectively, corresponding to the Zβ (F293-K366) domain. ADAR1 Zβ is coloured in dark blue. ZBP1 Zα1 and Zα2 are coloured in pink and purple, respectively.

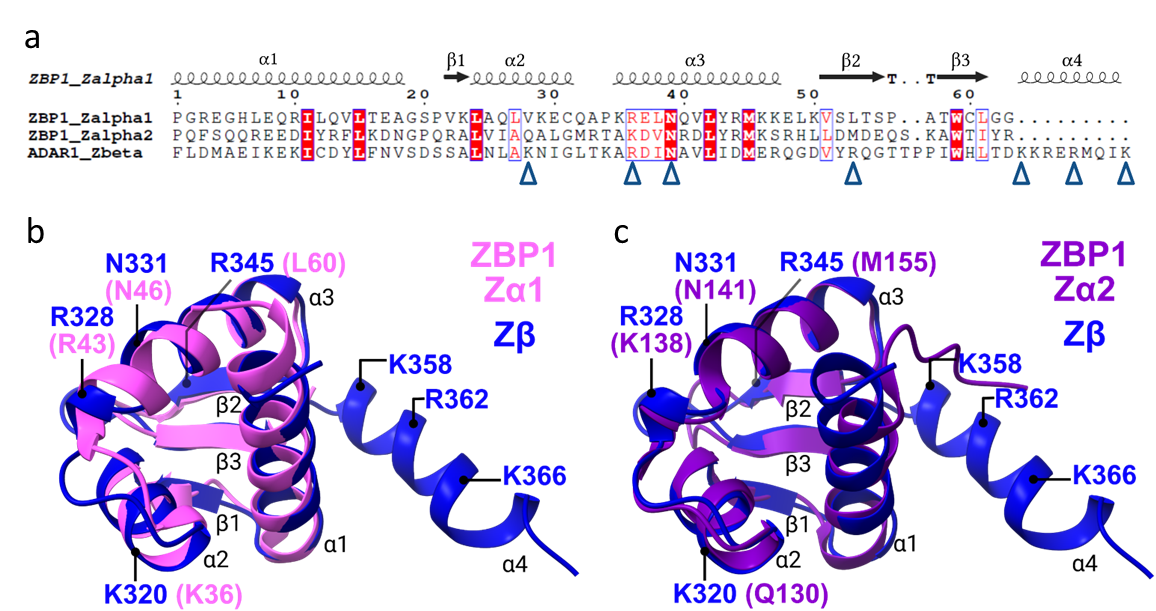

**Figure S7.** Kinetics of interaction between TNPO1 and ADAR1 constructs using biolayer interferometry. Representative runs of **a.** dsRBD construct containing dsRBD3 domain, **b.** Zα domain, **c.** Zβ domain (wild type), **d.** Zβ domain (K320A/R328A), **e.** Zβ domain (K320A/N331A), **f.** Zβ domain (K320A/K358A), and **g.** Zβ domain (K320A/R362A), are shown. Binding affinity is listed for constructs that showed binding to TNPO1.

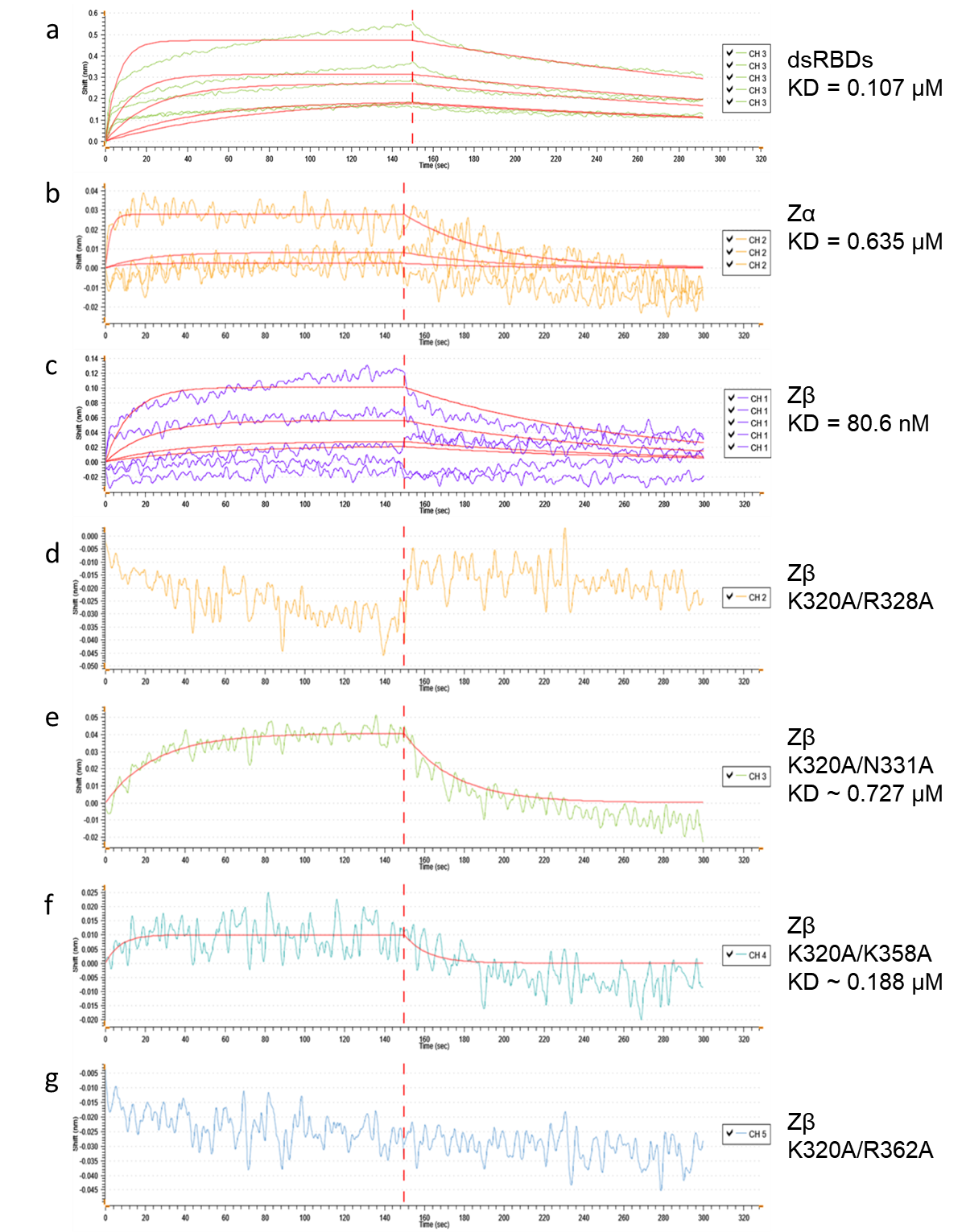

Tables

Table S1. Collection, refinement and validation statistics for cryo-EM density map and model. Collection details for the cryo-EM map presented in the text are shown above, and validation statistics for the model generated from the map are shown below.

|  | **EMDB: 100045** |
| --- | --- |
|  | **PDB: 34NN** |
| **Data collection and processing** |  |
| Detector | Falcon 4i |
| Magnification | 165,000 × |
| Voltage (kV) | 300 |
| Electron exposure (e–/Å^2^) | 50 |
| Defocus range (µm) | -0.5 ~-1.5 |
| Pixel size (Å) | 0.76 |
| Symmetry imposed | C1 |
| Micrographs collected (no.) | 5,000 |
| Initial particle images (no.) | 1,084,497 |
| Final particle images (no.) | 56,431 |
| Map resolution (Å) | 3.15 |
| FSC threshold micrographs | 0.143 |
| Map resolution range (Å) | 2.71-3.65 |
| **Refinement** |  |
| Initial model used (PDB code) | 2QMR, 1XMK, 2MDR |
| Model resolution (Å) | 3.2 |
| FSC threshold | 0.143-0.5 |
| Model resolution range (Å) | 1.6-3.1 |
| Map sharpening B-factor (Å^2^) | 77.9 |
| **Model composition** |  |
| Non-hydrogen atoms | 7882 |
| Protein residues | Protein: 993 |
| Nucleotide | 0 |
| Water | 1 |
| Ligands | 0 |
| **B factors (Å^2^)** |  |
| Protein | 27.75/166.77/73.92 |
| Nucleotide | N/A |
| Water | 63.98/63.98/63.98 |
| Ligand | N/A |
| **R.M.S. Deviations** |  |
| Bonds length (Å) | 0.003 (0) |
| Bonds Angle (˚) | 0.562 (0) |
| **Validation** |  |
| MolProbity score | 2.24 |
| Clashscore | 5.30 |
| Rotamer outliers (%) | 4.78 |
| **Ramachandran plot** |  |
| Favored (%) | 93.18 |
| Allowed (%) | 6.61 |
| Disallowed (%) | 0.20 |

Table S2. Comparison of buried solvent-accessible surface areas at the interfaces of TNPO1 with ADAR1 Zβ, dsRBD3, and representative PY-NLS-containing cargoes. Buried solvent-accessible surface areas (BSAs) were calculated using the UCSF ChimeraX “measure buriedarea” command and are reported in Å².

| **PDB / Interface** | **Buried Surface Area (Å^2^)** |
| --- | --- |
| Zβ-TNPO1 | 1356.7 |
| dsRBD3-TNPO1 | 1106.5 |
| 2OT8 | 894.03 |
| 5TQC | 488.18 |
| 8SGH | 466.33 |
| 4OO6 | 443.71 |
| 4FDD | 1097.5 |
| 5YVH | 1013.8 |
